# EHMT2 Inactivation in Pancreatic Epithelial Cells Shapes the Transcriptional Landscape and Inflammation Response of the Whole Pancreas

**DOI:** 10.1101/2024.03.14.584700

**Authors:** Gareth Pollin, Angela J. Mathison, Thiago M. de Assuncao, Anju Thomas, Lida Zeighami, Ann Salmonson, Hongfei Liu, Guillermo Urrutia, Pallavi Vankayala, Stephen J Pandol, Michael T. Zimmermann, Juan Iovanna, Victor X. Jin, Raul Urrutia, Gwen Lomberk

## Abstract

The Euchromatic Histone Methyl Transferase Protein 2 (EHMT2), also known as G9a, deposits transcriptionally repressive chromatin marks that play pivotal roles in the maturation and homeostasis of multiple organs. Recently, we have shown that EHMT2 inactivation alters growth and immune gene expression networks, antagonizing KRAS-mediated pancreatic cancer initiation and promotion. Here, we elucidate the essential role of EHMT2 in maintaining a transcriptional landscape that protects organs from inflammation. Comparative RNA-seq studies between normal postnatal and young adult pancreatic tissue from *EHMT2* conditional knockout animals (*EHMT2^fl/fl^*) targeted to the exocrine pancreatic epithelial cells (*Pdx1-Cre* and *P48^Cre/+^*), reveal alterations in gene expression networks in the whole organ related to injury-inflammation-repair, suggesting an increased predisposition to damage. Thus, we induced an inflammation repair response in the *EHMT2^fl/fl^*pancreas and used a data science-based approach to integrate RNA-seq-derived pathways and networks, deconvolution digital cytology, and spatial transcriptomics. We also analyzed the tissue response to damage at the morphological, biochemical, and molecular pathology levels. The *EHMT2^fl/fl^* pancreas displays an enhanced injury-inflammation-repair response, offering insights into fundamental molecular and cellular mechanisms involved in this process. More importantly, these data show that conditional EHMT2 inactivation in exocrine cells reprograms the local environment to recruit mesenchymal and immunological cells needed to mount an increased inflammatory response. Mechanistically, this response is an enhanced injury-inflammation-repair reaction with a small contribution of specific EHMT2-regulated transcripts. Thus, this new knowledge extends the mechanisms underlying the role of the EHMT2-mediated pathway in suppressing pancreatic cancer initiation and modulating inflammatory pancreatic diseases.

## INTRODUCTION

The pancreas arises from a small cluster of cells in the embryonic gut tube during development (Joglekar, Parekh, and Hardikar 2007). This process is tightly regulated by a complex network of signaling pathways and transcription factors, which control pancreatic progenitor cell specification, proliferation, and differentiation (Jarc et al. 2023). This glandular organ comprises different cell types, including endocrine cells that produce hormones and exocrine cells that produce digestive enzymes (Overton and Mastracci 2022). The proper development and function of the pancreatic cells are critical for maintaining health and preventing diseases, such as diabetes and pancreatitis (Karpińska and Czauderna 2022). These conditions are associated with defects in the molecular pathways that regulate pancreas development and function (Thrower, Husain, and Gorelick 2008; Polireddy and Chen 2016). However, there is limited knowledge on the precise mechanisms underlying these defects, and further research is still needed to better understand the complex molecular interactions within the pancreas.

The pathogenesis of acute pancreatitis is multifaceted and involves both local and systemic inflammatory responses. Epigenetic regulatory mechanisms play an important role in controlling the inflammatory cascade (Zhou et al. 2022). Various studies have investigated the role of epigenetic modifications in regulating inflammation and acute pancreatitis (Sandoval et al. 2016; Sun et al. 2021). DNA methylation is one of the most extensively studied epigenetic modifications in the context of acute pancreatitis. Studies have shown altered DNA methylation levels on genes involved in inflammation and oxidative stress in the pancreas during acute pancreatitis episodes(Natale et al. 2019; Sun et al. 2021). For instance, promoter hypermethylation of anti-inflammatory genes such as *IL10* and *SOCS3* has been associated with decreased expression and increased inflammation in the pancreas (Yin et al. 2015). Similarly, hypomethylation of pro-inflammatory *NFKB1* promoter regions increases expression and enhances pancreas inflammation. Histone modifications have also been implicated in the pathogenesis of acute pancreatitis. For example, histone H4 acetylation in the IL-1β promoter region correlates with increased expression of this pro-inflammatory cytokine in the pancreas (Pedersen et al. 2022). However, further research is needed to fully understand the complex interplay between epigenetic regulators and the inflammatory cascade in acute pancreatitis, which could lead to the development of new therapies for this debilitating disease.

Euchromatic histone-lysine N-methyltransferase 2 (EHMT2/G9a) is a methyltransferase that catalyzes the mono-and dimethylation of lysine 9 on histone H3 (H3K9me1/2), leading to transcriptional repression. EHMT2 has been implicated in regulating various cellular processes beyond gene expression, including cellular differentiation and DNA repair (Jan et al. 2021). While EHMT2 directly regulates the differentiation and function of various immune cell types, including T cells, B cells, and macrophages, several studies also support the role of EHMT2 in non-immune cells to control inflammatory responses (Mourits et al. 2021; Scheer and Zaph 2017). For instance, EHMT2 represses the expression of pro-inflammatory cytokines, such as TNF, in tumor cells to promote breast cancer recurrence (Mabe et al. 2020). Studies on vascular smooth muscle cells have also implicated EHMT2 in attenuating the IL-6 inflammatory response in atherosclerotic lesions (Harman et al. 2019). Furthermore, liver-specific EHMT2 knockout leads to an enhanced proinflammatory response in lipopolysaccharide (LPS)-induced liver injury model (Lu, Lei, and Zhang 2019). Recently, we have shown that EHMT2 inactivation antagonizes oncogenic KRAS-mediated pancreatic cancer initiation and promotion by altering growth and immune gene expression networks (Urrutia et al. 2021). EHMT2 knockout driven by either *Pdx1-Cre* or *P48^Cre/+^* demonstrated that this pathway is not required for pancreas exocrine development and is tolerated in this organ under basal contexts. However, the impact of EHMT2 on the transcriptome during early development and under the inflammatory stressor of acute pancreatitis remains unknown.

Here, we investigate the role of EHMT2 in postnatal murine pancreas development and caerulein-induced acute pancreatitis. EHMT2 inactivation results in distinct transcriptional landscapes during pancreatic maturation, suggesting an increased susceptibility of this organ to an injury-inflammation-repair process. Congruently, in response to the induction of acute pancreatitis, we show that EHMT2 inactivation leads to a more aggressive inflammatory response. Thus, this study advances our understanding of the epigenetic regulation and molecular mechanisms that play a role in pancreatic development, homeostasis, and diseases.

## RESULTS

### LOSS OF EHMT2 INCREASES THE PROPENSITY OF THE NORMAL PANCREAS TO INJURY-INFLAMMATION

Previously, we reported that mice with pancreas-specific *EHMT2* knockout develop and grow normally, adopting the right size, shape, and histology (Urrutia et al. 2021). While this phenotype indicates that this protein is not essential for organ development, it does underscore a function for EHMT2 in regulating gene expression networks during pancreatic maturation and pancreatitis. We harvested pancreas from *Pdx1-Cre; Ehmt2^fl/fl^* (*Ehmt2^fl/fl^*) and *Pdx1-Cre;Ehmt2^+/+^*(*Ehmt2^+/+^*) mice at both postnatal (PN) day 10 and young adulthood (YA) 4-5 weeks of age and performed RNA-seq. Principal component analysis (PCA) shows a defined clustering of PN and YA *Ehmt2^fl/fl^* animals apart from their control groups (Figure 1A). We identified 143 and 125 unique differentially expressed genes (DEGs) between *Ehmt2^fl/fl^* and *Ehmt2^+/+^* for YA and PN, respectively, with 26 overlapping between both age groups (Figure 1B). Using these DEGs, we carried out the deconvolution analysis through Quantiseq given that these mice were not immuno-stimulated in order to focus on the immune composition based on specific gene markers. This analysis revealed alterations in the relative proportions of immune cell populations induced by EHMT2 inactivation at different developmental times. During the YA stage, we found that the mature pancreas was less susceptible to baseline changes of immune cells with EHMT2 inactivation, resulting in a 7.5% decrease in marker genes for dendritic cells as well as a 7.3% increase in neutrophils and a minor 0.2%, increase in NK cells. In contrast, the PN *Ehmt2^fl/fl^* pancreas had a 55.3% increase in dendritic cell populations along with 42.1% and 13.2% decreases in neutrophil and NK cells, respectively (Figure 1C). We performed network enrichments to infer the function of the transcriptional landscape of the pancreas from these animals (Figure 1D). Upregulated genes showed significant enrichment for fibrin clot formation, hypoxia, and KRAS signaling up in the PN *Ehmt2^fl/fl^* mice compared to those with *Ehmt2* intact. The YA group was also enriched for these pathways and others, including erythrocyte gas exchange, extracellular matrix formation, and the scavenger receptor pathway. In contrast, no significant pathway enrichment was found among the downregulated DEGs. Comparison of molecular signatures in the YA upregulated DEGs group using MSigDB revealed enrichment of growth inhibitory pathways like P53 combined with KRAS signaling up, a phenomenon known to result in replication stress, cell cycle arrest, and apoptosis. More specifically we found upregulation of key kinases involved in the p21 pathway, such as *p21*, *Chek2* and *Ccng1* (Supplementary file), which we previously show causes cell-cycle arrest in the presence of activated KRAS in acinar cells (Urrutia et al. 2021). Moreover, these changes were accompanied by hypoxia, angiogenesis, and epithelial-mesenchymal transition (Figure 1D). Thus, we found that loss of *Ehmt2* most notably results in de-repression or upregulation of genes, in particular those that relate to development of an injury-inflammation-repair pathway.

**Figure 1:**
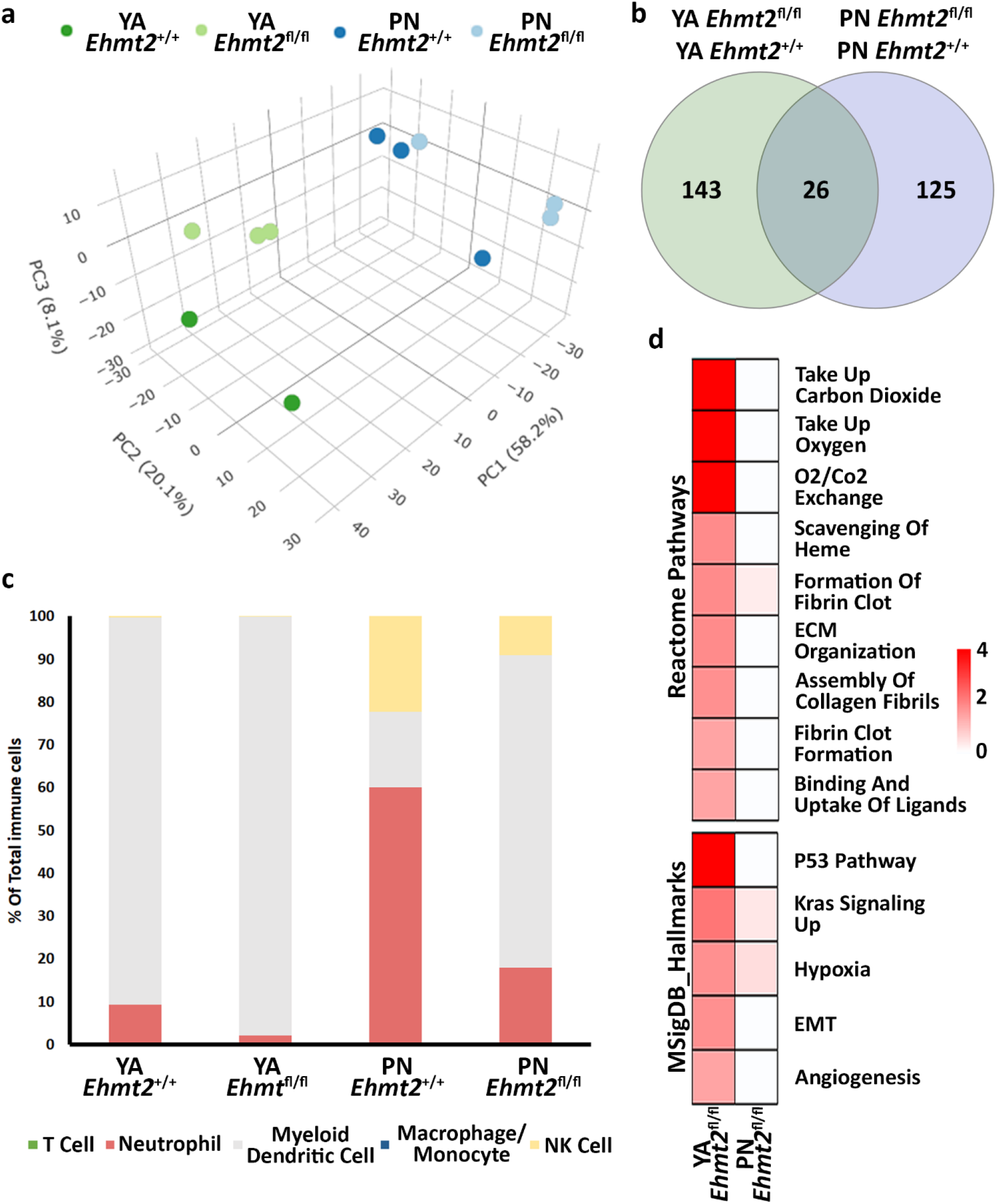
*Ehmt2* inactivation modulates the epigenetic landscape of mouse pancreas towards stress-related pathways. (**a**) PCA based on DEGs from RNA-seq conducted on pancreas tissue from Perinatal (PN) and young adult (YA) mice with and without *Ehmt2* is shown. (**b**) Venn diagram illustrates the variation in DEGs upon *Ehmt2* knockout (*Ehmt2^fl/fl^*) in the pancreas compared to *Ehmt2* wild-type (*Ehmt^+/+^*) mice for YA and PN cohorts. (**c**) Quantiseq deconvolution analysis predicts immune cell composition based on the percentage of total immune cells. (**d**) MSigDB Hallmarks and Reactome pathway enrichment analyses reveal the activation of functional pathways for differentially expressed genes in *Ehmt2^fl^*^/fl^ animals compared to their respective *Ehmt2*^+/+^ counterparts.

Next, we performed integrative analysis and functional modeling, by using the natural language processing-based identification of gene sets relationships from the ShinyGO suite combined with semantic-based algorithms using the DEGs from the across the different transcriptional landscapes, as input (Ge, Jung, and Yao 2020). This approach yielded several functional gene groups associated with pancreatic maturation in the *Ehmt2^fl/fl^* pancreas. For instance, we found higher representation of genes related to multiple diverse cellular functions including adhesion, extracellular matrix organization (ECM), inflammation response, stress and apoptosis, digestive enzymes metabolism, transport, epigenetics, and cell motility (Table 1). We also noted a differential activation of the hemoglobin locus control region (LCR) for Hba-a1, Hba-a2, Hbb-bs, and Hbb-bt, which serves as a pathognomonic indicator of alterations in the 3D organization of nuclei upon Ehmt2 loss. Furthermore, transcription factor motif analysis identified key upstream regulators of all DEGs related to EHMT2 inactivation, among which we find main transcriptional regulators of several pathways congruent with the mSigDB functional enrichments (enrichment of transcription factor binding motifs in DEG promoters; Table 2). In summary, loss of Ehmt2 results in dysregulation of the epigenetic landscape affecting essential signaling pathways, such as p53 and KRAS, which are known to cause replication stress, arrest cell proliferation, and induce apoptosis, combined with changes in the immune cell populations, suggesting an important function for Ehmt2 in maintaining pancreatic homeostasis and a susceptibility of *Ehmt2^fl/fl^* mice to heightened injury-inflammation repair responses.

**Table 1:**
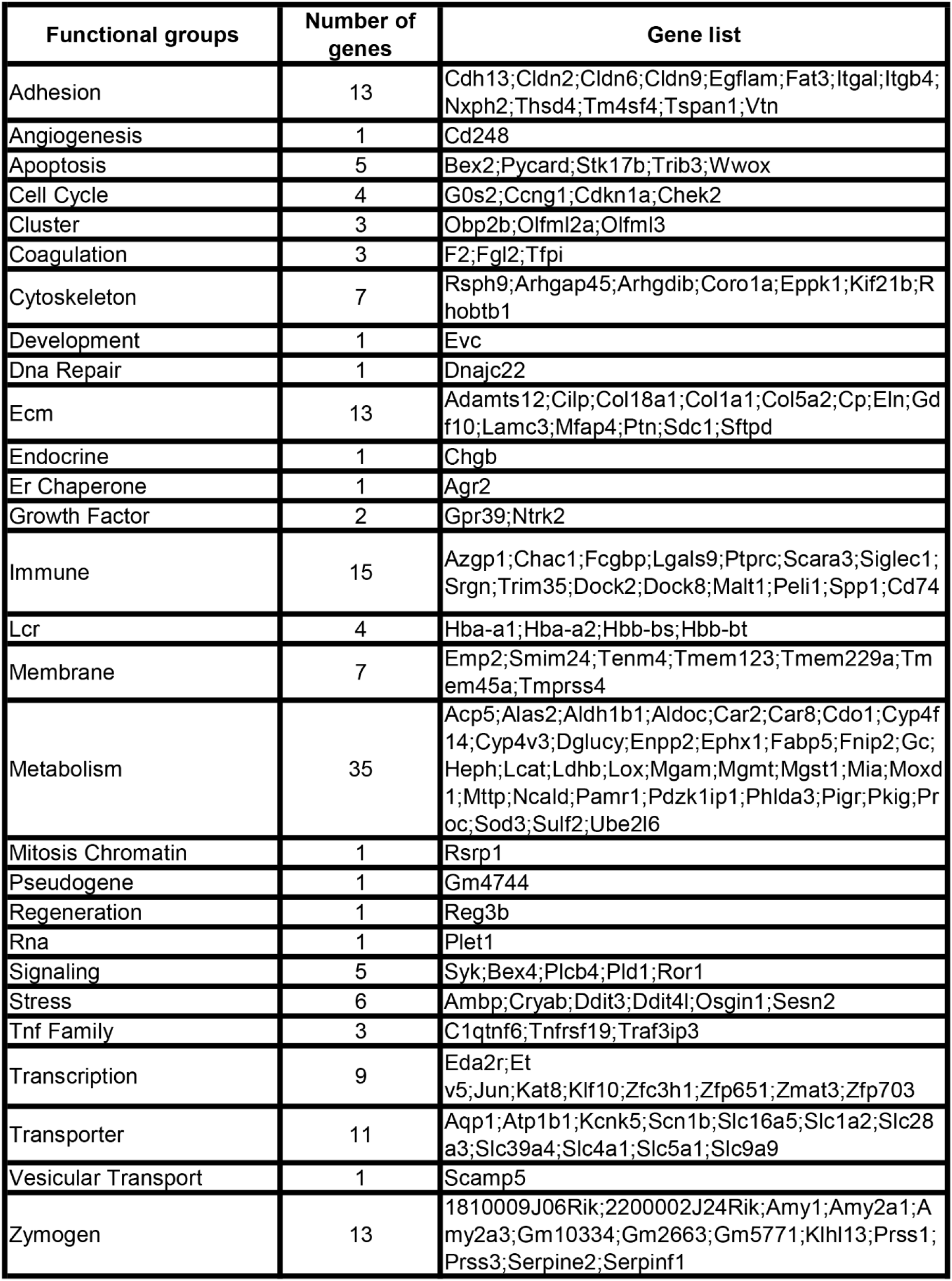
Functional annotation of DEGs regulated by *Ehmt2* inactivation in young adult mice.

**Table 2:**
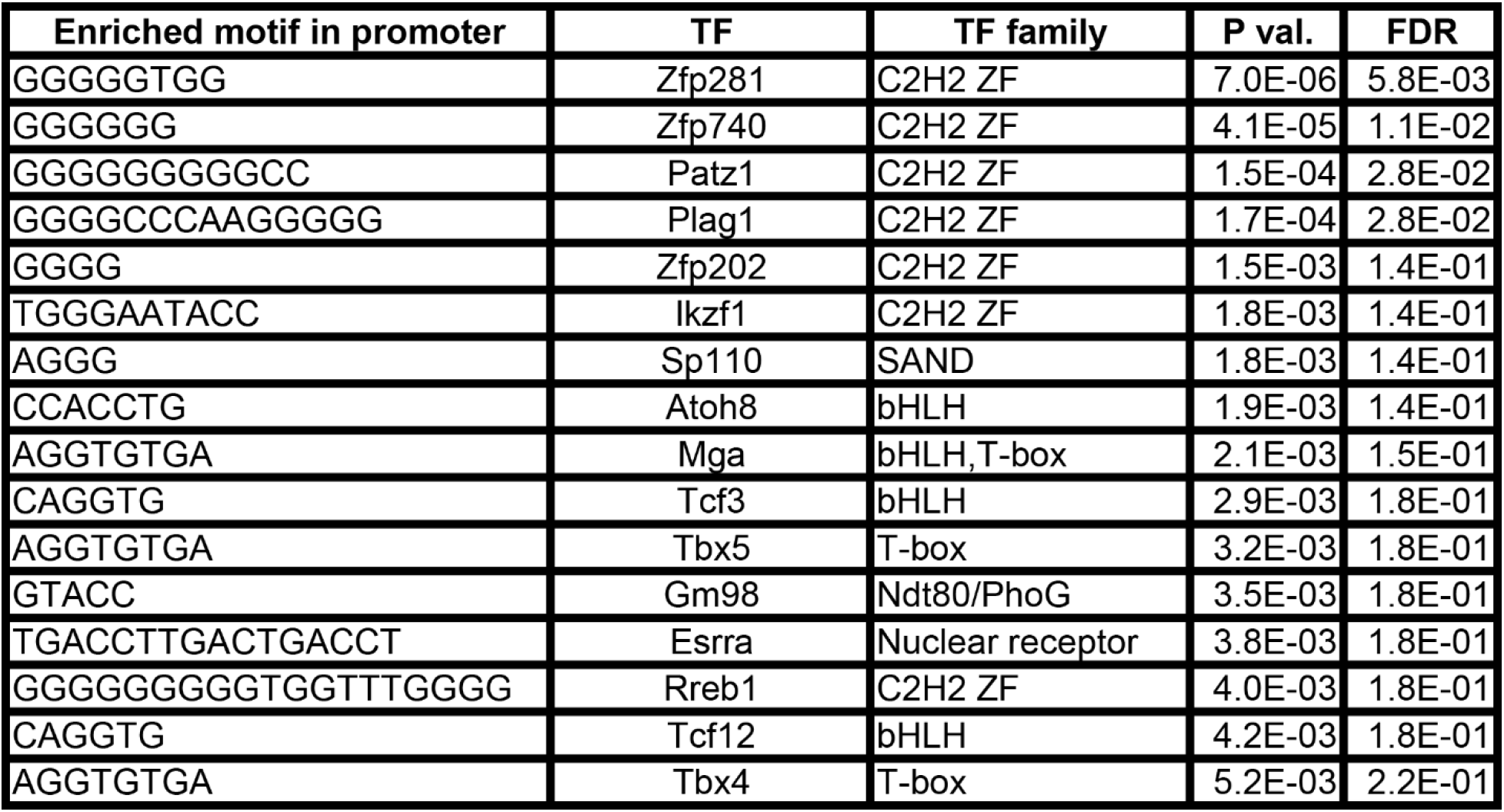
Transcriptional factor enrichment of DEGs upon *Ehmt2* inactivation in young adult mice.

### EHMT2 INACTIVATION IN ACINAR PANCREATIC CELLS ENHANCES THE RESPONSE TO ORGAN INFLAMMATION

To evaluate the relative contribution of Ehmt2 to the epithelial cell response in pancreatic inflammation, we repeatedly injected *Ehmt2^+/+^*and *Ehmt2^fl/fl^* mice with caerulein to induce acute pancreatitis and performed RNA-seq. PCA analysis showed a clear separation of mice subjected to caerulein compared to their untreated counterparts (Figure 2A). Notably, the untreated samples, regardless of *Ehmt2* inactivation status, clustered closely together on one end of the PCA plot. In contrast, samples undergoing acute pancreatitis displayed a distinct separation between the *Ehmt2^+/+^*and *Ehmt2^fl/fl^* pancreatic samples (Figure 2A). Using pairwise analysis between caerulein and untreated groups, we found 5,263 DEGs. Of these DEGs, 1,812 were shared between *Ehmt2^+/+^* and *Ehmt2^fl/fl^* with acute pancreatitis, consisting of 764 upregulated and 1,048 downregulated DEGs (Figure 2B-C). In terms of unique DEGs, 1,838 were distinctly upregulated upon acute inflammation in the *Ehmt2^fl/fl^* mice versus 189 in the *Ehmt2^+/+^*group (Figure 2B). Correspondingly, 1,283 were exclusively downregulated during acute pancreatitis in the *Ehmt2^fl/fl^* animals compared to 142 in the *Ehmt2^+/+^* (Figure 2C). Interestingly, a large subset of the genes that were found in the *Ehmt2^+/+^* animals, which would constitute the response to acute pancreatitis, overlapped with the *Ehmt2^fl/fl^* mice (80% of upregulated and 91% of downregulated genes). Moreover, 70% of upregulated genes and 55% of downregulated genes that were differentially regulated in *Ehmt2^fl/fl^* animals were unique to the loss of *Ehmt2^fl/fl^*. This highlights that Ehmt2 plays a major role during acute pancreatitis in maintaining transcriptional homeostasis in response to injury. Through heatmap clustering analyses of these DEGs, we identified that the genes differentially regulated in both *Ehmt2^+/+^* and *Ehmt2^fl/fl^*animals presented with an amplified response when *Ehmt2* was inactivated (Figure 2D). This finding in the *Ehmt2^fl/fl^* mice with acute pancreatitis is congruent with a principal role of Ehmt2 in mediating gene expression in response to tissue injury. We found that E*hmt2^+/+^* and *Ehmt2^fl/fl^*mice with acute pancreatitis, compared to their untreated counterparts, exhibited significant upregulation of genes that were enriched in injury-inflammatory pathways through activation of the oncogenic KRAS, P53, TNFα and TGF-β signaling pathways (Figure 2E). Examination of upstream regulatory elements that were found to have an enhanced inflammatory response when *Ehmt2* was inactivated, revealed a network of genes that include *Il1b*, *Il1r1*, *Tnf*, *Csf3r* and members of the Ccl family, which are known recruit and activate immune cells. With Il1r1 serving as the primary receptor for Il1b, this pathway is recognized for its role in triggering NF-κB signaling pathway activation, which we also found (Table 3). We also detected increased gene expression networks related to immune cell activation and function, such as *Slamf* and cell surface molecules receptors. Similarly, the upregulation of chemokine signaling by C-X-C motif chemokine ligand families, indicate a heightened recruitment and migration of immune cells to inflamed pancreatic tissue. Notably, we also found upregulation of integrin and Interleukin cytokines, with roles in immunomodulatory processes that influence the balance between pro-inflammatory and anti-inflammatory responses. Thus, the increased expression of various inflammatory mediators highlights a cascade of cellular events that seem to culminate in the initiation and propagation of the enhanced immune response discovered in the *Ehmt2^fl/fl^* mice. In addition, there was increased representation of genes involved in growth factor signaling, cytoskeletal and cell adhesion, and regulation, DNA replication and repair, metabolic and oxidative stress, and RNA processing that collectively function to enhance cell survival, proliferation, and tissue regeneration during pancreatitis. Furthermore, we observe derepression of the Beta Globin Locus Control Region (LCR), a crucial regulatory element known for its involvement in intrachromosomal looping (Table 3). This event is a marker of 3D reorganization of the nucleus (Deng et al. 2012). Contrastingly, the downregulated genes showed substantial enrichment of metabolic and proteostasis pathways, such as fatty and bile acid metabolism and unfolded protein response. This observation highlights that acute pancreatitis causes drastic changes to the pancreas gene expression landscape reflecting that normal functionality of the pancreas is reduced (Figure 2E). Furthermore, we observed an increased enrichment of genes involved in a repair response, such as activation of mTORC1, PI3K-Akt, Androgen and Estrogen hormone signaling pathways. Lastly, we observed activation of genes involved in other signaling networks to help the pancreas tissue cope with damage caused by acute pancreatitis, such as angiogenesis to compensate for increased metabolic demand (Figure 2E). Notably, in the *Ehmt2^fl/fl^* pancreas tissues, the enrichment and overall number of genes for each ontology was found to be of greater significance with an increased number of genes in each of the categories, specifically for the UV response, cholesterol, MTORC1, myogenesis, estrogen, and reactive oxygen related pathways (Figure 2E). Thus, this data further highlights the EHMT2-mediated functions that are necessary for the injury-inflammation-repair response during acute pancreatitis.

**Figure 2:**
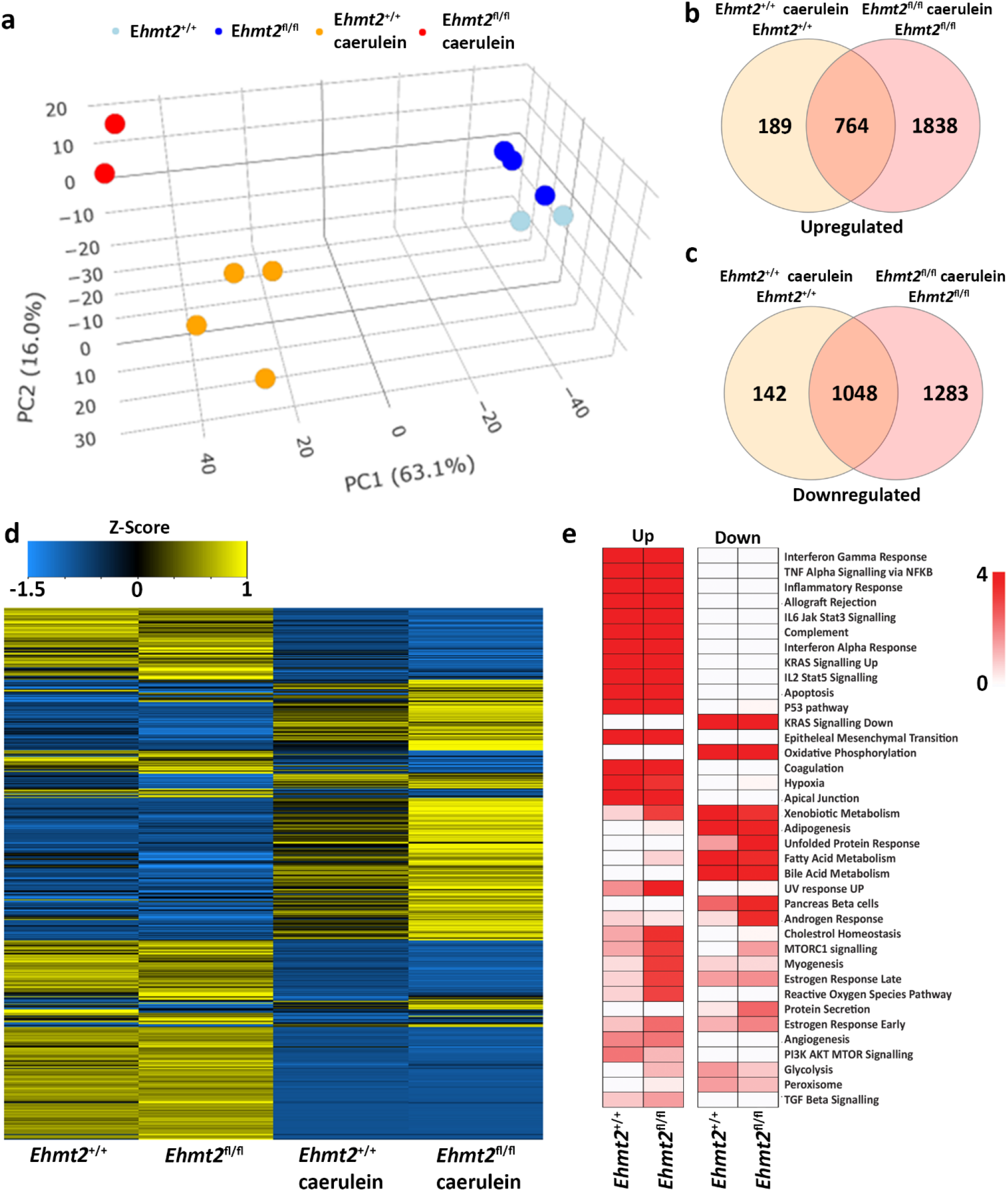
Ehmt2 is crucial for maintaining faithful transcriptional homeostasis in response to injury during acute pancreatitis. (**a**) PCA based on DEGs from RNA-seq conducted on pancreas tissue from *Ehmt2^+/+^* and *Ehmt2^fl/fl^* adult mice both with and without induction of acute pancreatitis is shown. Venn diagram illustrates the overlap in significant DEGs between mice treated with caerulein and untreated mice, comparing *Ehmt2^+/+^* and *Ehmt2^fl/fl^* animals for both (**b**) upregulated and (**c**) downregulated genes. (**d**) Heatmap displays the average expression of DEGs that were significant in at least one condition for *Ehmt2^+/+^*and *Ehmt2^fl/fl^* animals in absence or presence of caerulein treatment. (**e**) MSigDB Hallmarks pathway enrichment analysis of significant DEGs between mice treated with caerulein and untreated mice in both *Ehmt2^+/+^* and *Ehmt2^fl/fl^* animals is shown for upregulated and downregulated genes.

**Table 3:**
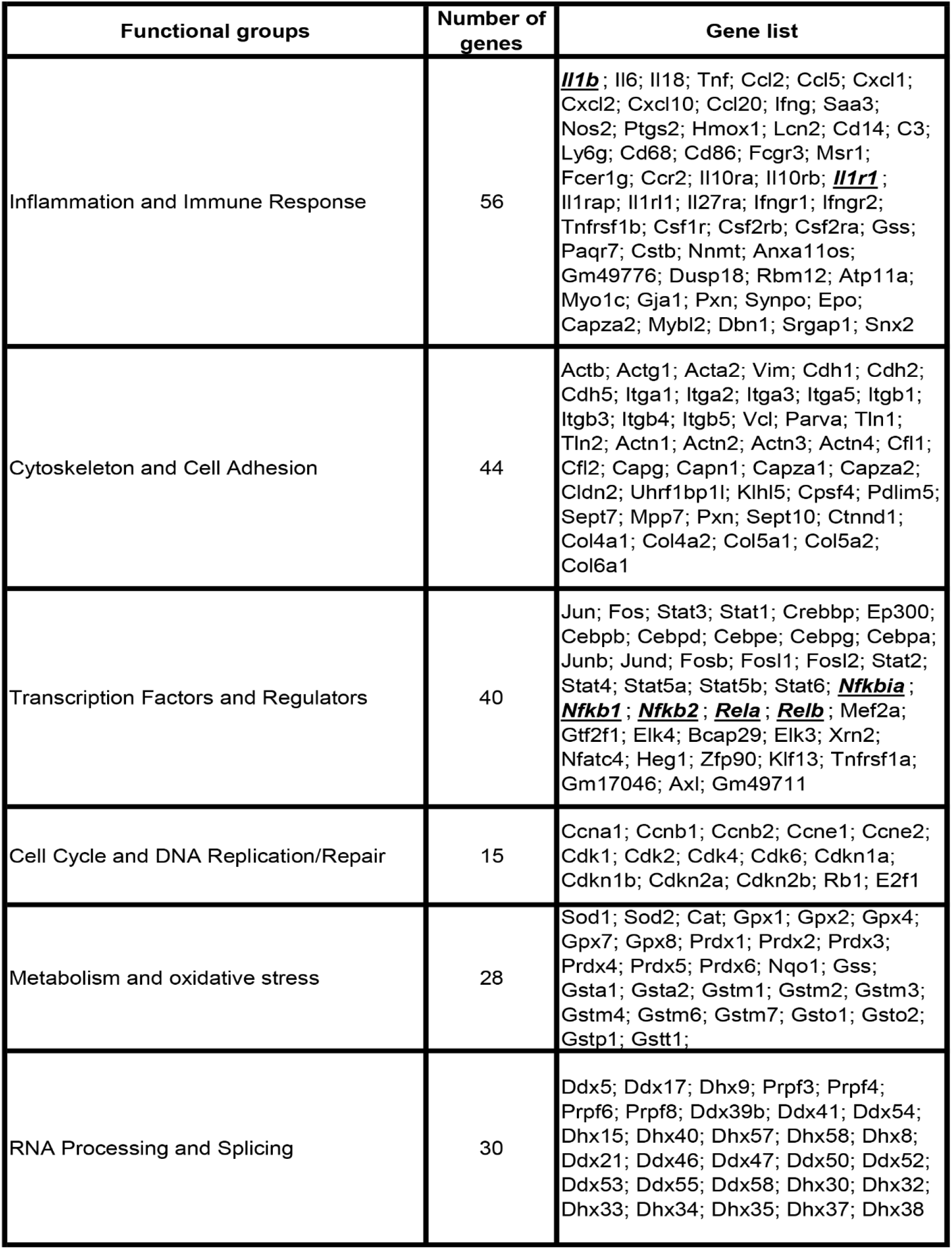
Enhanced expression/de-repression of Ehmt2-regulated genes that are associated with the inflammatory and immune response in acute pancreatitis.

To reveal differences in the composition of the immune cell infiltrate across the total cells found between the *Ehmt2^+/+^* and *Ehmt2^fl/fl^* pancreas of caerulein-treated animals, we used the MCP Counter, an expanded deconvolution method due to the increased inflammation under these conditions (Figure 3A). The induction of acute pancreatitis highlighted immune cell patterns that distinguished *Ehmt2^+/+^* from *Ehmt2^fl/fl^*, also revealing an enhanced immune response when Ehmt2 function is lost with the total number of immune cells increased by a total of 16.6%. Specifically, we identified an increase of 1.9% in T-cells in animals with *Ehmt2* inactivation. Similar increases were found for canonical markers of neutrophils and myeloid dendritic cells with an increase of 1.9% and 1.2%, respectfully, for *Ehmt2^fl/fl^* animals compared to *Ehmt2^+/+^*mice. Notably, macrophages represented the most substantial increase of 11.6% upon *Ehmt2* knockout. (Figure 3A). These findings highlight the important role of Ehmt2 in regulating immune cell infiltration during acute pancreatitis. Thus, we used spatial transcriptomics to validate our findings from bulk RNA-seq. Analytically, we applied Seurat’s functionalities to define spatially variable genes (SVG) across tissue samples. To deconvolute the cell types within the spatial transcriptomic spots, we employed marker genes obtained from Zhou et al (2022) and CellMarker (Zhang et al. 2019). We calculated the average number of spots containing SVGs indicative of immune infiltration by T cells (*Cd3e*), neutrophils (*S100a8*), dendritic cells (*Ccr7*), macrophages (*Apoe*), and NK cells (*Nk7g*) for *Ehmt2^+/+^* and *Ehmt2^fl/fl^* mice. In concordance with our bulk RNA-seq data, we detected the enhanced immune response in *Ehmt2^fl/fl^* compared to the *Ehmt2^+/+^*mice with comparable distributions of specific immune subtypes (Figure 3A). An average of 7865 spots for *Ehmt2^+/+^* and 7542 for *Ehmt2^fl/fl^* mice were assigned in total across the tissue, and from these total spots, 0.8% contained SVG expression indicative of T-cell infiltrate in the *Ehmt2^+/+^* pancreas, which increased to 1.9% in *Ehmt2^fl/fl^*. Similarly, we found higher proportions of neutrophils and dendritic cells in *Ehmt2^fl/fl^* tissue (2.6% and 2.0%, respectively) compared to *Ehmt2^+/+^* (0.5% and 0.6%, respectively). Once again, the macrophage population signified the most striking change between *Ehmt2^+/+^* tissue (10.0%) and *Ehmt2^fl/fl^* (27.4%). Conversely, the difference in the proportions of NK cells were modest with 0.3% in *Ehmt2^+/+^* to 0.8% in *Ehmt2^fl/fl^* (Figure 3A). To more comprehensively address the complex pathology of pancreatic disease, we expanded our spatial transcriptomic analysis to other relevant cell types using previously reported markers as labeled in Figure 3B (Olaniru et al. 2023). We found a significant reduction in spots associated with the SVG marker for acinar cells (*Prss2*) in *Ehmt2^fl/fl^* samples, with only 66.6% of spots indicative of acinar cells (Figure 3B). In contrast, *Ehmt2^+/+^* tissues displayed a more typical distribution, with 94.4% of spots representing the presence of acinar cells. Similarly, *Ehmt2^fl/fl^* exhibited a decrease in duct cells compared to *Ehmt2^+/+^* (39.1% *vs.* 30.0%). On the other hand, features for islets (*Ins2*) (15.9% vs 17.4%), fibroblasts (*Cola1a*) (10.6% vs 12.7%), stellate cells (*Mmp14*) (1.1% vs 2.3%), and tuft cells (*Dclk1*) (1.0% vs 2.2%) showed slight increases when comparing *Ehmt^+l+^* tissues to *Ehmt2^fl/fl^*. Lastly, we evaluated SVGs that are hallmarks of epithelial-mesenchymal transition (EMT). Focusing on the gene *Spp1*, we found 25.7% of spots positive for EMT-like SVG expression in *Ehmt2^+/+^* mice with acute pancreatitis, which notably increased to 45.7% in *Ehmt2^fl/fl^*pancreas (Figure 3B). As the macrophage marker showed the highest increase across the immune cells, we correspondingly found a clear increase of high expressing *Apoe* spots, suggesting a higher infiltrate and immune response when *Ehmt2* is inactivated (Figure 3C). Thus, the results from spatial transcriptomics fundamentally confirmed our findings in bulk RNA-seq that Ehmt2 loss in the acinar cell triggers an increased immune infiltrate upon induction of acute pancreatitis.

**Figure 3:**
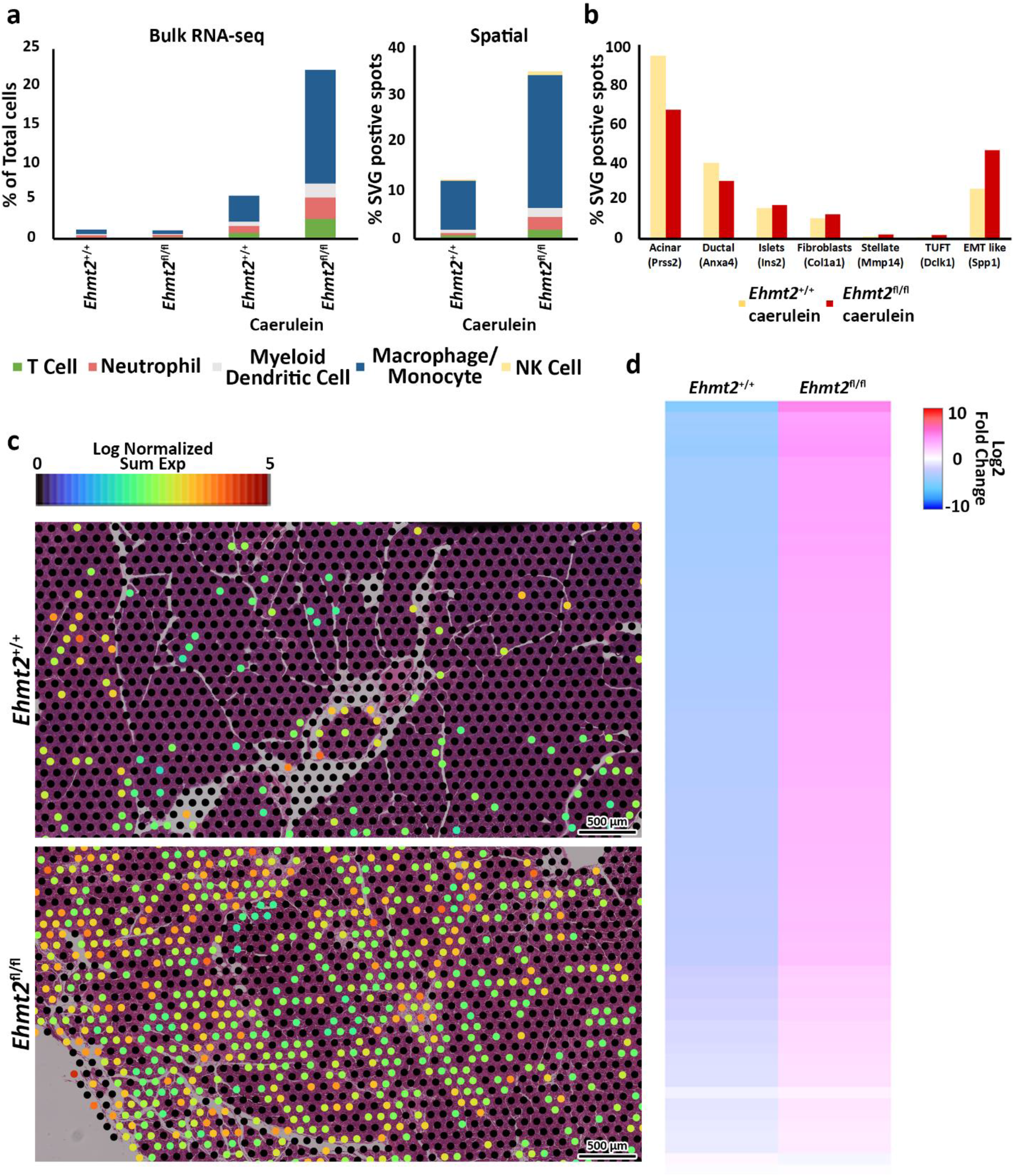
Ehmt2 deficiency promotes immune cell infiltration in injured pancreatic tissue. (**a**) MCPcounter deconvolution analysis predicts immune cell composition in the total cell population using Bulk RNA-seq data from *Ehmt2^+/+^*and *Ehmt2^fl/fl^* animals, in the absence or presence of caerulein treatment. Manual spatial deconvolution is conducted using spots positive for SVGs and stringent immune cell markers in pancreas samples from caerulein-treated *Ehmt2^+/+^* and *Ehmt2^fl/fl^* animals. (**b**) Bar graph shows pancreas cell deconvolution utilizing spots positive for SVGs of markers as indicated for a subset of pancreatic cells in pancreas tissue from caerulein-treated *Ehmt2^+/+^*and *Ehmt2^fl/fl^* animals. (**c**) Visualization of macrophage SVG-positive spots is overlayed on caerulein-treated *Ehmt2^+/+^* and *Ehmt2^fl/fl^* pancreas samples. (**d**) Heatmap displays log2 fold change for 11 genes within the HBB LCR control region on chromosome 7.

We further used the spatial analysis to investigate the normalized sum expression of genes within the HBB cluster which was observed to be dysregulated in the bulk RNA-seq, including Hbb-bs, Hbb-bt, Hbb-y, Hbb-hb2, Hbb-bh1, Olfr66, Olfr64, Olfr65, Olfr67, Olfr68, and Olfr69. Our analysis revealed that *Ehmt2^+/+^*mice exhibited an average of 7.51% of SVG spots positive for this group of genes, while *Ehmt2^fl/fl^* mice showed an increase to 13.54%. The mean expression level of these spots in *Ehmt2^+/+^* animals measured 0.15, whereas in the *Ehmt2^fl/fl^* samples, the mean expression significantly rose to 0.35 (Figure 3D). This looping mechanism plays a crucial role in controlling gene expression by altering active and inactive compartments of topologically associating domains in 3D. These collective findings suggest that Ehmt2 plays a crucial role in preserving the epithelial characteristics of both acinar and ductal cells in response to pancreatic injury, particularly during instances such as acute pancreatitis. This involvement is underscored by its role in gene repression and the regulation of 3D nuclear localization, emphasizing the multifaceted mechanisms through which Ehmt2 contributes to the maintenance of cellular identity in the context of physiopathological stimuli.

### EHMT2-MEDIATED PATHWAYS DURING PANCREAS INFLAMMATION ARE MODEL-INDEPENDENT

To confirm the enhanced tissue damage response during acute pancreatitis observed with loss of *Ehmt2*, we performed a pairwise analysis of the pancreatitis transcriptional landscape of the *Ehmt2* knockout generated either with the *Pdx1-Cre* (*Pdx1-Ehmt2^fl/fl^*) or *P48^Cre/+^* (*P48-Ehmt2^fl/fl^*) - driven models. This analysis will allow for the identification of model-independent genes involved in the response to injury during acute pancreatitis that are transcriptionally regulated by Ehmt2. PCA demonstrated that Cre animals with intact *Ehmt2* clustered together during the acute pancreatitis treatment regardless of the model, *Pdx1* or *P48^Cre/+^*, and separated from their *Ehmt2^fl/fl^* counterparts (Figure 4A). Analysis of both models together reveals 765 DEGs with the *P48-Ehmt2^fl/fl^* and 954 DEGs with the *Pdx1-Ehmt2^fl/fl^* model when compared to their *Ehmt2^+/+^* counterparts (Figure 4B-C). Across DEGs, 496 genes were shared among *Ehmt2^fl/fl^*mice with acute pancreatitis, regardless of the Cre-driver used (Figure 4B-C). More specifically, we found 650 upregulated DEGs in the *Pdx1-Ehmt2^fl/fl^* and 518 with the *P48-EHMT2^fl/fl^*, of which 347 DEGs were the same between the two models (Figure 4B). For downregulated DEGs, there were 139 DEGs in common between both models from the 304 downregulated DEGs in *Pdx1-Ehmt2^fl/fl^* and 257 in the *P48-EHMT2^fl/fl^*tissues (Figure 4C). The high reproducibility and separation between acute pancreatitis-induced animals with *Ehmt2* intact and those with *Ehmt2* loss, regardless of the Cre model, present in the PCA (Figure 4A) are also evident in the heatmap (Figure 4D). Ontological analysis of DEGs revealed significant enrichment of distinct functional groups, including inflammatory response and immune signaling, cell signaling and proliferation, as well as metabolism and hormonal regulation. For instance, upregulated DEGs in both the *Pdx1-Ehmt2^fl/fl^* and *P48-EHMT2^fl/fl^* acute pancreatitis groups, that were classified within the inflammatory response and immune signaling functional group, were enriched in pathways such as TNF-α signaling via NF-κB, interferon gamma response, allograft rejection, IL-6/JAK/STAT3, and IL-2/STAT5 signaling. For the cell signaling and proliferation functional group, we found enrichment in KRAS signaling up, EMT, angiogenesis, the p53 pathway, and apoptosis, cholesterol homeostasis, xenobiotic metabolism, estrogen response late, and estrogen response early were enriched under metabolism and hormonal regulation DEGs for the *Pdx1-Ehmt2^fl/fl^*mice. Similar enrichment was observed for upregulated DEGs in *P48-EHMT2^fl/fl^*mice. Conversely, downregulated DEGs in the *Pdx1-Ehmt2^fl/fl^* and the *P48* model with acute pancreatitis model were only significantly enriched for networks that signal KRAS down (Figure 4E). Thus, whether *Ehmt2* is knocked out of the pancreas using either the *Pdx1-Cre* or *P48^Cre/+^*model, the modified transcriptional impact of acute pancreatitis compared to their counterparts with *Ehmt2* intact are similar, highlighting that Ehmt2 participates primarily in preventing an enhanced injury-inflammatory repair response in this organ.

**Figure 4:**
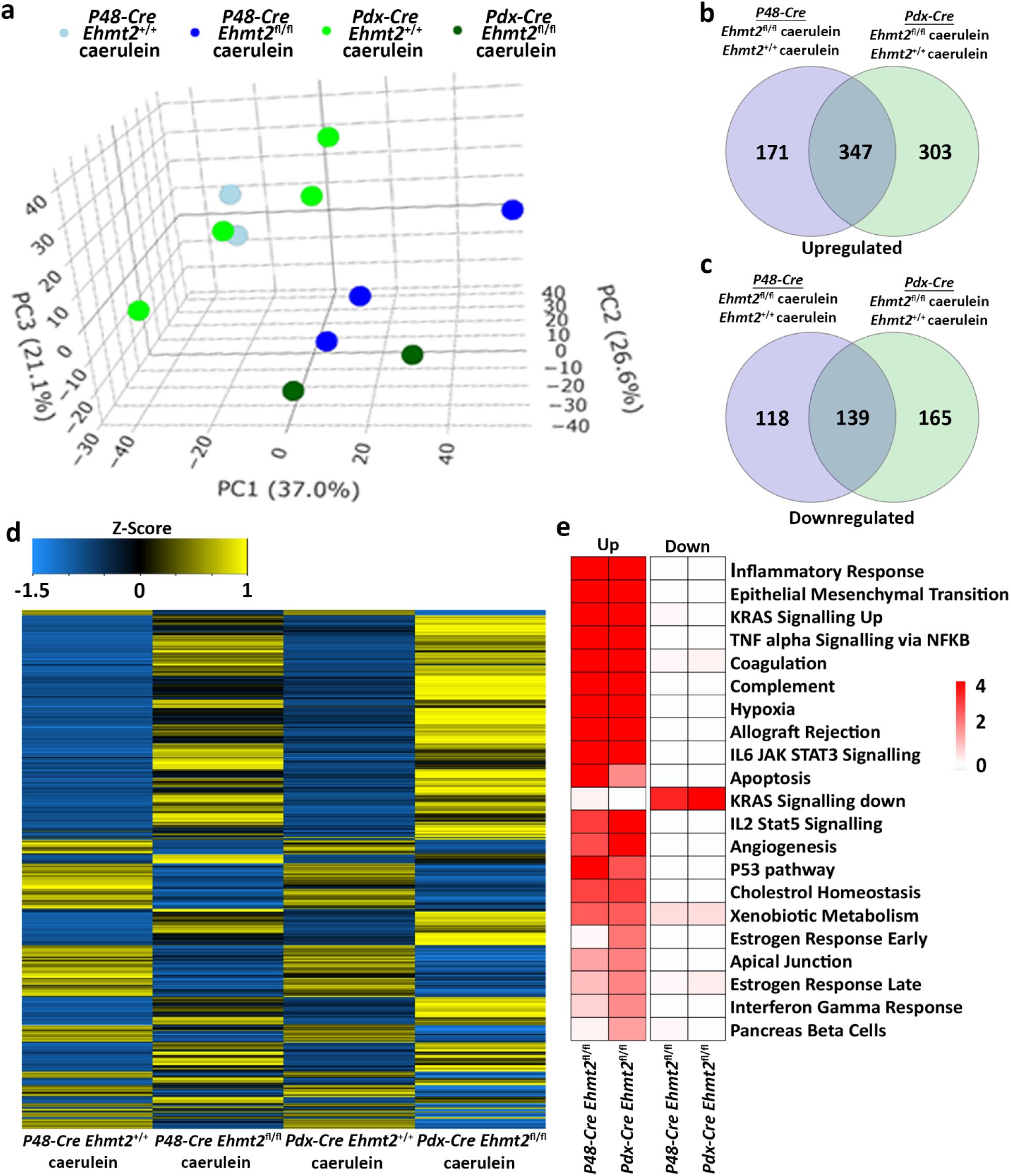
Ehmt2-mediated transcriptional regulation during acute pancreatitis is model independent. (**a**) PCA based on DEGs from RNA-seq conducted on pancreas tissue from adult *Ehmt2^+/+^* and *Ehmt2^fl/fl^* mice with acute pancreatitis for both *Pdx-Cre* and *P48-Cre* driven models is shown. Venn diagram illustrates the overlap in significant DEGs between pancreatitis-induced *Pdx-Cre* and *P48-Cre* models with *Ehmt2* inactivation compared to their *Ehmt2^+/+^* counterparts for (**b**) upregulated and (**c**) downregulated genes. (**d**) Heatmap displays the average expression of DEGs that were significant in at least one condition for *Ehmt2^+/+^* and *Ehmt2^fl/fl^* animals treated with caerulein in both *Pdx-Cre* and *P48-Cre* models. (**e**) MSigDB Hallmarks pathway enrichment analysis is shown for significant DEGs between caerulein-treated *Pdx-Cre* and *P48-Cre* models in both *Ehmt2^+/+^* and *Ehmt2^fl/fl^* animals for both upregulated and downregulated genes.

Lastly, we performed further analysis using NLP and semantic-based algorithms on DEGs identified by the PDX and P48 models to determine shared pathways, which included signal transduction and immune defense (Table 4). We found enrichment in cell adhesion and extracellular matrix (ECM) genes, indicating changes in ECM organization, tissue remodeling processes, and immune cell trafficking within the pancreas during pancreatitis. Moreover, *Ehmt2^fl/fl^* animals upregulated several enzymes, transporters, and channels, which likely reflects the increased metabolic needs (Table 4). Additionally, we noted that derepression of *Il1b* and its known receptor *Il1r1*, as well as increased expression of upstream regulators involved in the Nfkb/Rel immune-mediated response were present in both the PDX and P48 models. Analysis of enriched cis-regulatory elements within promoter regions of these DEGs revealed associations with universal stripe factors such as ZFP281, PATZ1, SP, and KLF5 among others (Zhao et al. 2022). This suggests that Ehmt2 may play a role in the crosstalk mechanism between these universal stripe factors, facilitating chromatin accessibility (Table 5). Thus, the response to inflammation that we describe here for loss of Ehmt2 in the pancreas epithelial cell is primarily not mediated by tissue enriched Stripe Factors but rather general Stripe factors that drive responses in many organs, suggesting that these processes may have broader relevance.

**Table 4:**
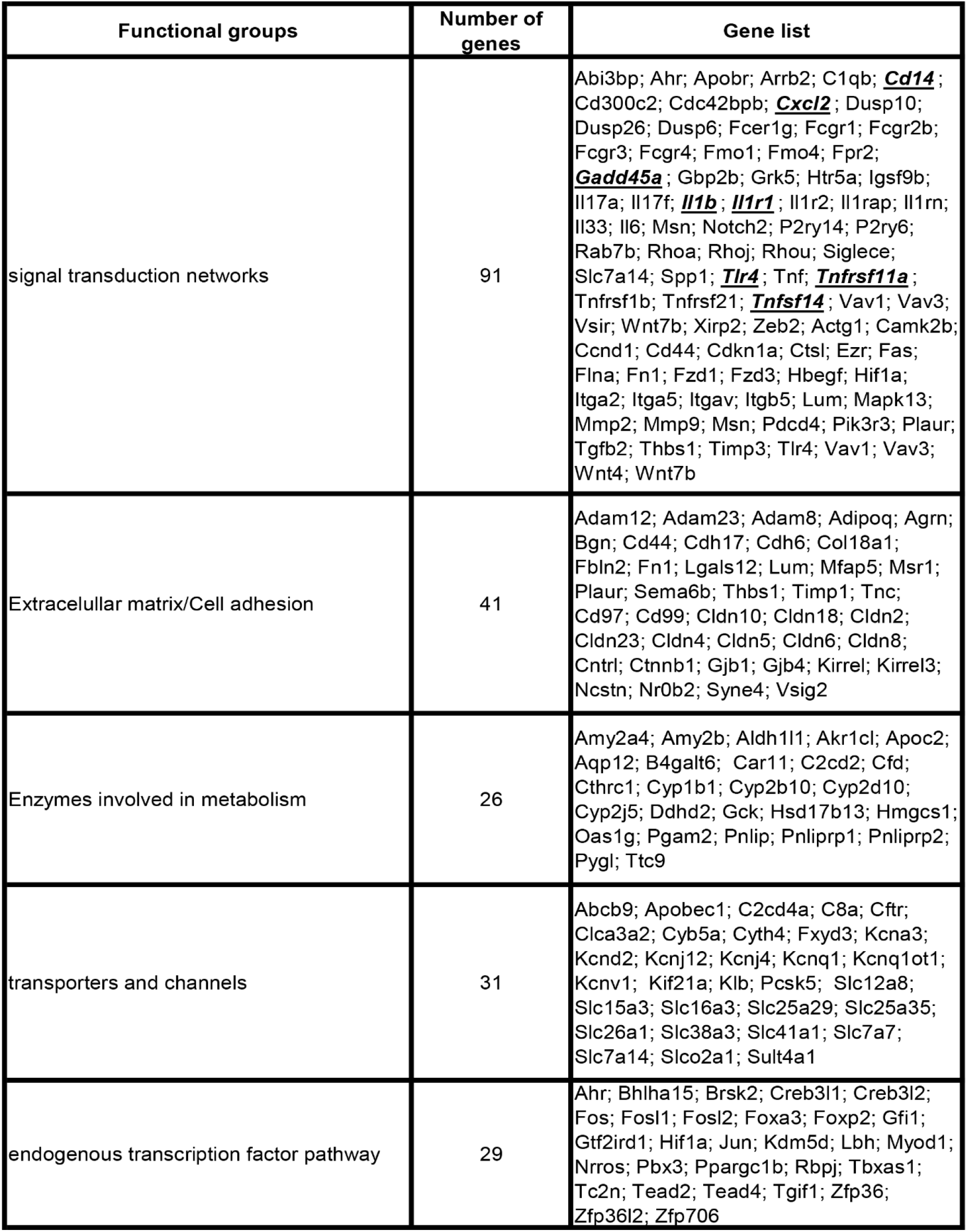
Integrative analysis by biological modeling of model-independent DEGs to determine shared Ehmt2-dependent pathways.

**Table 5:**
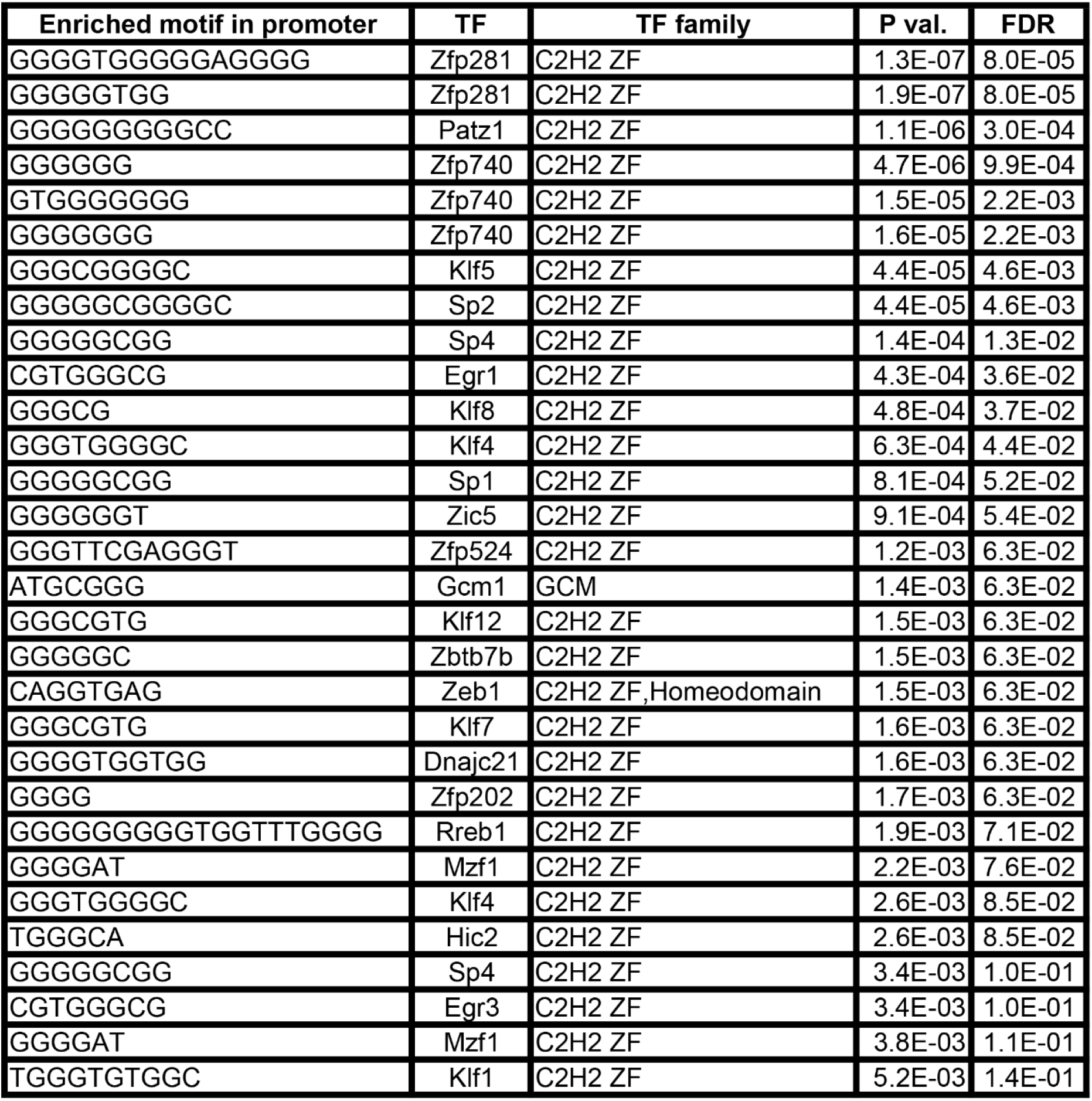
Transcriptional factor enrichment of DEGs by Ehmt2 during acute pancreatitis.

### MORPHOLOGICAL AND BIOCHEMICAL ASSESSMENT CONFIRMS THE IMPACT OF EHMT2 INACTIVATION ON PANCREAS INFLAMMATION IN BOTH THE PDX1- AND P48-DRIVEN MODELS

Biochemical investigation was performed on serum from both *Pdx1-Cre* and *P48^Cre/+^* caerulein-treated mice with or without *Ehmt2*. We found significantly decreased levels of albumin, total protein, blood urea nitrogen (BUN) and alkaline phosphatase (ALP) levels in *Ehmt2^fl/fl^* animals compared to *Ehmt2^+/+^* mice (Figure 5A).The decrease in circulating albumin, coupled with heightened inflammation in *Ehmt2^fl/fl^* mice, suggests a potential increase in capillary permeability and the efflux of circulating albumin into interstitial spaces (Soeters, Wolfe, and Shenkin 2019), which is seen in cases of acute pancreatitis (Xu et al. 2023). The reduction in albumin, along with diminished total protein levels in *Ehmt2^fl/fl^* serum implies a potential impairment in liver function (Soeters, Wolfe, and Shenkin 2019; Pompili et al. 2023; Wen et al. 2022; Rodriguez-Alvarez et al. 2023). Furthermore, decreased ALP levels in *Ehmt2^fl/fl^* serum may directly result from compromised liver function (Iluz-Freundlich et al. 2021). The alterations in serum constituents suggest that *Ehmt2* deficiency exacerbates inflammation under conditions of acute pancreatitis and may compromise the adjacent hepatic tissue. In line with these results, we found that the pancreas to body weight ratios increased when *Ehmt2* was inactivated during acute pancreatitis, likely resulting from a swollen and enlarged pancreas (Figure 5B). Indeed, histopathological examination of pancreatic tissue after induction of acute pancreatitis revealed that loss of *Ehmt2* from the pancreatic epithelium was characterized by an intensified reaction to caerulein administration with several notable histopathological changes compared to their *Ehmt2^+/+^* counterparts (Figure 5C). First, we detected a pronounced increase in inflammatory infiltrates, comprising a higher density of neutrophils, macrophages, and lymphocytes within the pancreatic parenchyma and interstitial spaces (Figure 5D). Additionally, we found more extensive edema with interlobular and intralobular spaces displaying marked expansion due to enhanced vascular permeability. Heightened vascular fragility and leakage were also evidenced by hemorrhagic areas in tissue from the *Ehmt2^fl/fl^* animals. Furthermore, acinar cells in *Ehmt2^fl/fl^* exhibited severe damage, with widespread cellular vacuolization, loss of cytoplasmic granularity, and cellular disintegration, leading to disrupted tissue architecture (Figure 5D). In contrast, acinar cells in *Ehmt2^+/+^* animals maintained a more intact structure even with the caerulein-induced inflammatory insult. The fibrosis was also notably heightened in *Ehmt2^fl/fl^* animals. Subsequently, we assessed whether this exacerbated response to acute pancreatitis in *Ehmt2^fl/fl^* animals was associated with acinar cell death. We found increased immunohistochemical staining for cleaved caspase 3 and TUNEL-positive cells, indicating increased apoptosis within the pancreatic tissue of *Ehmt2^fl/fl^* mice (Figure 5E-F). In summary, loss of *Ehmt2* in the pancreatic epithelium results in an augmented response to acute pancreatitis characterized by higher inflammatory cell infiltration, edema, acinar cell damage and death, as well as augmented necrosis.

**Figure 5:**
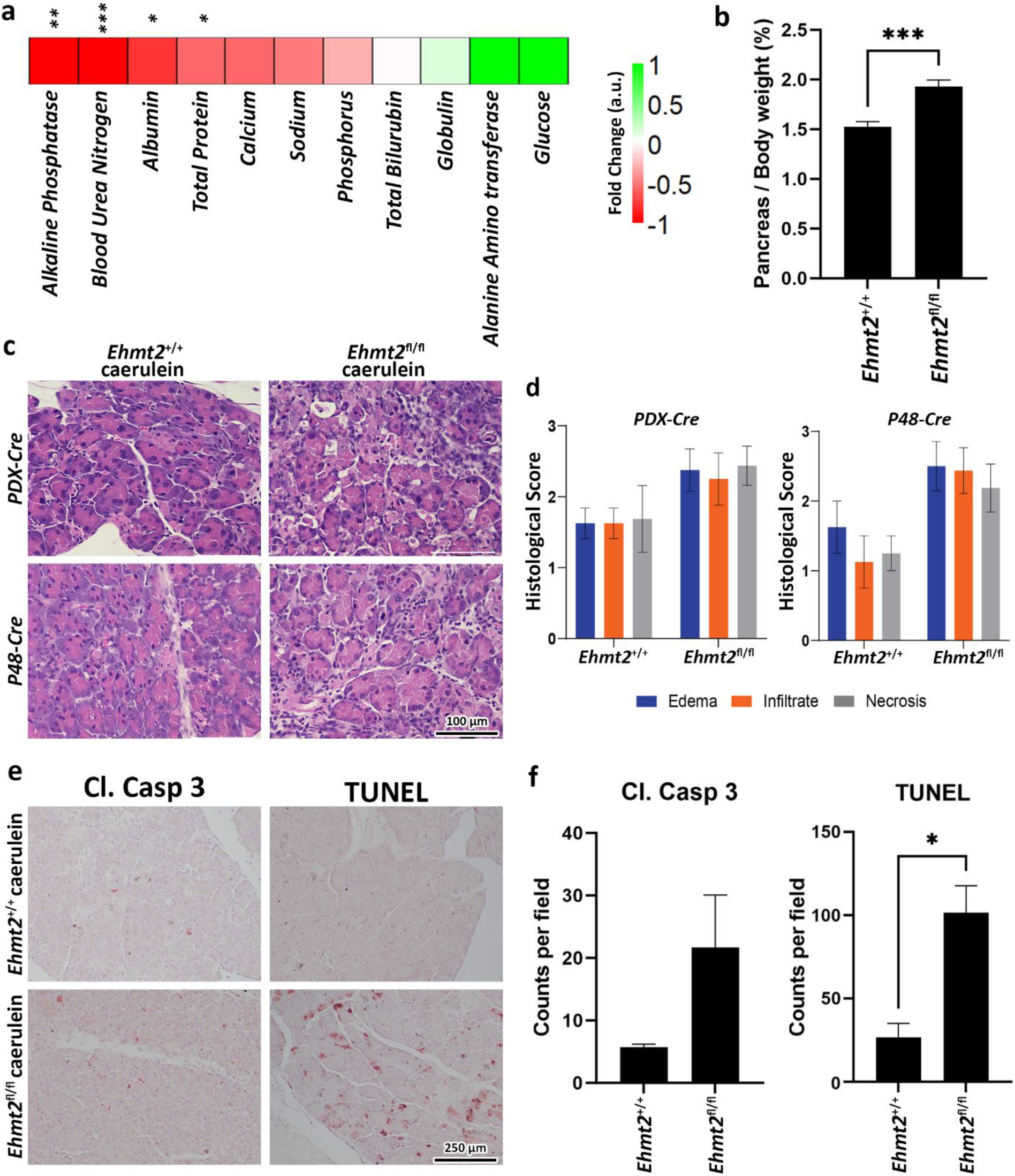
Ehmt2 deficiency in the pancreas epithelium increases tissue damage and deterioration in acute pancreatitis. (**a**) Heatmap shows the various circulatory markers in plasma of mice with acute pancreatitis. The difference in change between *Ehmt2^+/+^* and *Ehmt2^fl/fl^* groups have been depicted where +1 shows highest and -1 shows lowest difference in change. (**b**) Graph shows increased pancreas-to-body weight ratios after *Ehmt2* inactivation compared to control *Ehmt2^+/+^* animals in caerulein-induced acute pancreatitis. (**c**) Representative histology of caerulein-induced pancreatitis from *Pdx1-Cre* and *P48-Cre Ehmt2^+/+^* and *Ehmt2^fl/fl^* pancreas tissue is shown by hematoxylin-eosin-staining. Scale bar, 100 μm. (**d**) Pancreatitis histological scores for edema, infiltrate and necrosis were assessed from four random fields (20× objective) per slide, each containing at least 1,000 cells per field. Scores, expressed as mean ± SD scores, were calculated for each individual parameter. (**e**) Representative images are shown for immunohistochemical staining for cleaved Caspase 3 and TUNEL in pancreas tissue from *Pdx1-Cre* and *P48-Cre Ehmt2^+/+^* and *Ehmt2^fl/fl^* mice with caerulein-induced pancreatitis. (**f**) Bar graph depicts quantification for cleaved Caspase 3 and TUNEL assay reveals increased apoptosis in *Ehmt2^fl/fl^*compared to control *Ehmt2^+/+^* animals in caerulein induced pancreatitis. Plasma Samples; n=6. Pancreas/Body weight samples; n = 3. Caspase/TUNEL assay; n=6. (*, P < 0.05; **, P < 0.01; ***, P < 0.001; t-test)

To validate the aggravated inflammatory response in *Ehmt2^fl/fl^*animals in comparison to the *Ehmt2^+/+^*, we analyzed our spatial transcriptomics using a gene profile for the tumor inflammation signature (TIS) (Loh et al. 2023; Danaher et al. 2018). TIS expression in *Ehmt2^+/+^* was an average of 0.39, which increased to 0.65 in *Ehmt2^fl/fl^* pancreas tissues (Figure 6A). Overall, using a t-SNE plot, we found that 50.8% of SVG spots were TIS significant spots in the *Ehmt2^fl/fl^* tissues, and 45.3% were TIS significant in the *Ehmt2^+/+^* pancreas, of which most spots showed higher expression upon Ehmt2 knockout (Figure 6B). We found that all genes from the TIS were statistically significant between the *Ehmt2^fl/fl^* pancreas compared to the *Ehmt2^+/+^* controls during acute pancreatitis, with a range from 0.34 for *Psmb10* as the least log2 fold change to 3.48 for *Tigit* with the most significant change (Table 6). Moreover, the TIS-related SVGs were substantially increased in density when visualized overlaying the pancreas tissue for the *Ehmt2^fl/fl^* when compared to the *Ehmt2^+/+^* (Figure 6C). Using a gene signature for acute pancreatitis (Fang et al. 2022), we identified a marked increase of overall mean sum expression in *Ehmt2^fl/fl^* (3.3) animals in comparison to *Ehmt2^+/+^* (2.6) and an increase of 92% of positive SVGs for the *Ehmt2^fl/fl^*animals compared to 83% for the *Ehmt2^+/+^* mice (Figure 6D-E). Upon closer examination of individual genes, we observed that the genes in the signature for acute pancreatitis were predominantly overexpressed by a log2FC of >0.8 in the *Ehmt2^fl/fl^* group compared to *Ehmt2^+/+^*(Table 6). Using the spatial visualization tool, we observe substantial increased expression for each of the SVGs for the overall acute pancreatitis gene signature in the *Ehmt2^fl/fl^* pancreas compared to the *Ehmt2*^+/+^ organ, concordant with *Ehmt2* inactivation enhancing the inflammatory response in acute pancreatitis (Figure 6F). Only three genes (*Hspb1*, *Krt8* and *Hsp90aa1*) exhibited higher expression in the *Ehmt2^+/+^* pancreas, albeit with a log2FC no greater than 0.3. Thus, collectively, these results underscore the impact that *Ehmt2* inactivation within the pancreas epithelium has on the response to inflammatory injury of the whole organ.

**Figure 6:**
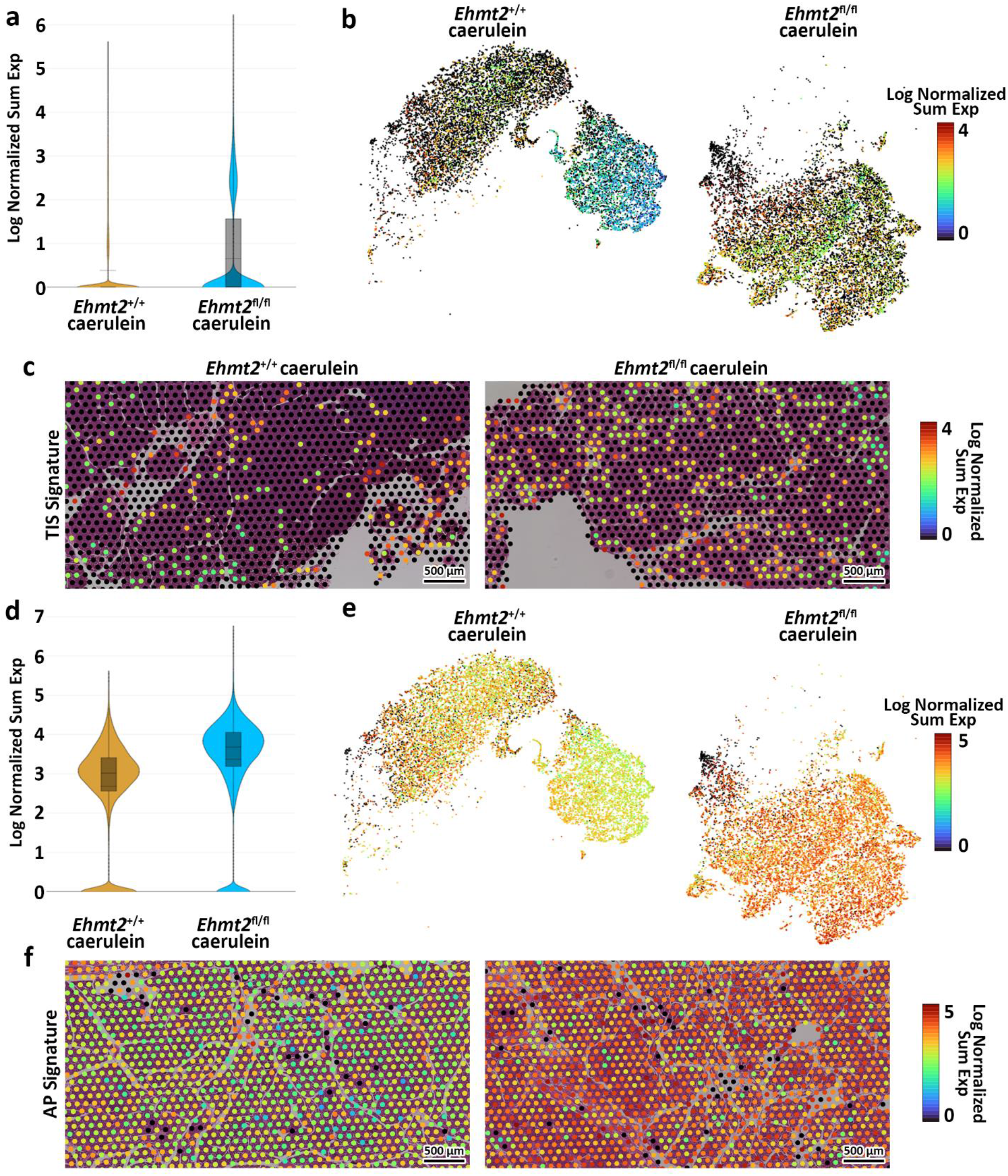
Loss of Ehmt2 results in derepression of tumor associated and acute pancreatitis inflammatory markers increasing susceptibility to tissue damage in the pancreas. (**a**) Violin plots display the log fold change of normalized gene expression, describing expression of the tumor inflammation signature (TIS) in spatial transcriptomics derived from pancreas tissues from acute pancreatitis induced *Ehmt2^+/+^* and *Ehmt2^fl/fl^* mice. (**b**) t-SNE plot illustrates the distribution of SVG positive spots corresponding to genes from the TIS. (**c**) Visualization overlays depict the spatial distribution and expression of TIS positive spots in acute pancreatitis samples from *Ehmt2^+/+^*and *Ehmt2^fl/fl^* mice. (**e**) Violin plots display the log fold change of normalized gene expression, describing expression of the acute pancreatitis (AP) signature in spatial transcriptomics derived from pancreas tissues of *Ehmt2^+/+^* and *Ehmt2^fl/fl^* mice with caerulein-induced acute pancreatitis. (b) t-SNE plot illustrates the distribution of SVG positive spots corresponding to genes from the AP signature. (c) Visualization overlays depict the spatial distribution and expression of AP signature positive spots in acute pancreatitis samples from *Ehmt2^+/+^*and *Ehmt2^fl/fl^* mice. Scale bars represent the log normalized sum expression of selected genes.

**Table 6:** Spatial transcriptomics shows increased expression of TIS and acute pancreatitis signature markers in tissues from caerulein-treated *Ehmt2^fl/fl^ animals*.

| FeatureID | Signature | FeatureName | WT Average | G9a KO Average | G9a KO Log2 FC | P-Value |
| --- | --- | --- | --- | --- | --- | --- |
| ENSMUSG00000031897 | Tumor Inflammation Signature | Psmb10 | 0.052 | 0.066 | 0.34 | 2.3E-02 |
| ENSMUSG00000026104 | Tumor Inflammation Signature | Stat1 | 0.006 | 0.010 | 0.67 | 1.0E-03 |
| ENSMUSG00000035042 | Tumor Inflammation Signature | Ccl5 | 0.001 | 0.002 | 0.80 | 3.3E-02 |
| ENSMUSG00000035914 | Tumor Inflammation Signature | Cd276 | 0.016 | 0.038 | 1.22 | 1.1E-14 |
| ENSMUSG00000042190 | Tumor Inflammation Signature | Cmklr1 | 0.005 | 0.032 | 2.64 | 5.9E-49 |
| ENSMUSG00000016498 | Tumor Inflammation Signature | Pdcd1lg2 | 0.003 | 0.018 | 2.79 | 2.4E-42 |
| ENSMUSG00000004612 | Tumor Inflammation Signature | Nkg7 | 0.001 | 0.008 | 2.89 | 3.4E-25 |
| ENSMUSG00000016496 | Tumor Inflammation Signature | Cd274 | 0.003 | 0.022 | 2.93 | 1.3E-47 |
| ENSMUSG00000035000 | Tumor Inflammation Signature | Dpp4 | 0.002 | 0.014 | 2.95 | 3.1E-35 |
| ENSMUSG00000030124 | Tumor Inflammation Signature | Lag3 | 0.003 | 0.021 | 3.02 | 4.6E-50 |
| ENSMUSG00000048521 | Tumor Inflammation Signature | Cxcr6 | 0.002 | 0.016 | 3.10 | 1.7E-41 |
| ENSMUSG00000029417 | Tumor Inflammation Signature | Cxcl9 | 0.003 | 0.025 | 3.12 | 3.8E-48 |
| ENSMUSG00000031551 | Tumor Inflammation Signature | Ido1 | 0.001 | 0.010 | 3.22 | 7.5E-27 |
| ENSMUSG00000053977 | Tumor Inflammation Signature | Cd8a | 0.001 | 0.013 | 3.36 | 3.1E-43 |
| ENSMUSG00000071552 | Tumor Inflammation Signature | Tigit | 0.001 | 0.009 | 3.48 | 3.1E-27 |
| ENSMUSG00000004951 | acute pancreatitis | Hspb1 | 0.206 | 0.165 | -0.32 | 2.2E-02 |
| ENSMUSG00000049382 | acute pancreatitis | Krt8 | 1.003 | 0.944 | -0.09 | 5.3E-01 |
| ENSMUSG00000021270 | acute pancreatitis | Hsp90aa1 | 0.036 | 0.034 | -0.06 | 7.5E-01 |
| ENSMUSG00000025283 | acute pancreatitis | Sat1 | 0.063 | 0.110 | 0.81 | 2.7E-09 |
| ENSMUSG00000115338 | acute pancreatitis | Pnp | 0.090 | 0.179 | 1.00 | 3.7E-14 |
| ENSMUSG00000030342 | acute pancreatitis | Cd9 | 0.035 | 0.074 | 1.08 | 5.3E-15 |
| ENSMUSG00000007041 | acute pancreatitis | Clic1 | 0.106 | 0.253 | 1.25 | 4.5E-22 |
| ENSMUSG00000009687 | acute pancreatitis | Fxyd5 | 0.032 | 0.096 | 1.56 | 3.0E-31 |
| ENSMUSG00000022146 | acute pancreatitis | Osmr | 0.009 | 0.029 | 1.76 | 7.4E-26 |
| ENSMUSG00000032231 | acute pancreatitis | Anxa2 | 0.073 | 0.313 | 2.11 | 4.8E-62 |
| ENSMUSG00000021091 | acute pancreatitis | Serpina3n | 0.016 | 0.079 | 2.31 | 6.0E-58 |
| ENSMUSG00000026628 | acute pancreatitis | Atf3 | 0.014 | 0.072 | 2.41 | 2.7E-62 |
| ENSMUSG00000028494 | acute pancreatitis | Plin2 | 0.018 | 0.130 | 2.83 | 1.7E-94 |
| ENSMUSG00000005087 | acute pancreatitis | Cd44 | 0.065 | 0.506 | 2.97 | 2.9E-125 |
| ENSMUSG00000005413 | acute pancreatitis | Hmox1 | 0.018 | 0.142 | 2.98 | 2.2E-105 |
| ENSMUSG00000026822 | acute pancreatitis | Lcn2 | 0.019 | 0.251 | 3.74 | 1.8E-171 |
| ENSMUSG00000019122 | acute pancreatitis | Ccl9 | 0.025 | 0.471 | 4.21 | 8.0E-243 |

## DISCUSSION

The current study offers significant insights into the role of EHMT2 in pancreatic development and its response to inflammatory stimuli. Clinical and experimental observations have shown that individual responses to pancreatitis-causing stimuli vary, suggesting contributions from both genetic factors and environmental exposures (Weiss, Laemmerhirt, and Lerch 2021). However, the specific mechanisms underlying these individual responses, particularly concerning chromatin proteins that epigenetically regulate transcriptional landscapes, remain poorly understood. Thus, the current study was designed to directly investigate the role of Ehmt2, a robust transcriptional repressor, in experimentally induced acute pancreatitis. Utilizing a systems biology approach and genetically engineered mouse models, we employed RNA-Seq with deconvolution into a digital cytology approach, as well as spatial transcriptomics, to define the emergent properties resulting from the genetic inactivation of *Ehmt2* in pancreatic acinar cells. Our results reveal, for the first time, that *Ehmt2* inactivation increases the propensity of the normal pancreas to injury-inflammation, suggesting its crucial role in maintaining pancreatic homeostasis and moderating inflammatory responses. Notably, we observed significant transcriptional differences in *Ehmt2^fl/fl^* mice during postnatal and young adult stages, indicating the involvement of this epigenomic regulator in fine-tuning gene expression networks essential for pancreatic maturation. Furthermore, the induction of acute pancreatitis in *Ehmt2^fl/fl^* mice resulted in a more aggressive inflammatory reaction, highlighting its role in suppressing inflammatory gene networks and restraining the pancreatic inflammatory response. This enhanced response was consistently confirmed by morphological and biochemical evidence across different models, including both *Pdx1-Cre* and *P48^Cre/+^*-driven models, indicating its robustness and independence from specific genetic backgrounds. These findings provide important insights into the molecular mechanisms underlying pancreatic diseases, particularly those with an inflammatory component, shedding light on the pathogenesis of these conditions.

The observation of enhanced inflammatory response in *Ehmt2^fl/fl^*mice holds significant implications from an evolutionary standpoint. The *Ehmt2* gene itself is embedded within a cluster of inflammatory genes (Scheer and Zaph 2017), suggesting a potential evolutionary pressure that has preserved its role as a key regulator of inflammatory pathways across successive generations. This arrangement implies a functional importance of EHMT2 in modulating inflammatory responses, with its genetic proximity to other inflammatory genes possibly indicating a coordinated regulatory mechanism. This evolutionary perspective underscores the critical role of EHMT2 in maintaining homeostasis and regulating inflammatory processes within the pancreas and potentially other tissues.

Mechanistically, we identify key transcriptional nodes in epithelial cells that alter the profiles of secreted factors, impacting the reactions and function of surrounding cells. *Ehmt2* inactivation leads to an enhanced inflammatory response rather than a distinct one, evidenced by changes in the levels of chemokine family members crucial for leukocyte recruitment to inflammatory sites (Popiolek-Barczyk et al. 2020). This observation has broad implications for various diseases, including developmental disorders, inflammatory conditions, and cancer. Moreover, we found the increased expression of several inflammatory mediators, such as *Il1b*, *Il1r1*, *TNF* among others, that likely augment the activation of the NF-κB signaling pathway. Interestingly, EHMT2 has been previously linked to the NF-κB signaling pathway, including its interaction with the NF-κB transcription factor RelB (Harman et al. 2019; Chen et al. 2009). These findings underscore the intricate interplay among EHMT2, inflammatory mediators, and the NF-κB signaling pathway, shedding light on the molecular mechanisms underlying inflammatory responses across various disease contexts. In particular, our findings regarding acute pancreatitis offer valuable insights into how dysregulation of epithelial cells can drive inflammatory responses and influence disease progression within the pancreas, while also potentially informing our understanding of related conditions, such as chronic pancreatitis, autoimmune pancreatitis, and pancreatic cancer.

We also report that EHMT2 functions as a critical regulator of transcriptional programs essential for maintaining pancreatic acinar cell function. These cells must cyclically produce large amounts of enzymes for macromolecule digestion, necessitating tight coordination of nuclear functions, RNA and ribosome production in the nucleolus, mRNA splicing in the spliceosome, protein synthesis in abundant rough ER, and cytoplasmic ribonucleoprotein-rich granules for storage (Logsdon and Ji 2013). Conditional *Ehmt2* inactivation in acinar cells leads to dysregulation of genes influencing the way the entire organ reacts to inflammation. This dysregulation occurs via two major cellular mechanisms: increased activity of gene networks antagonizing the cell cycle, and alterations in this transcriptional regulator that modify expression of molecular mediators, exacerbating acute pancreatitis severity. Furthermore, our findings suggest that *Ehmt2* inactivation also affects nuclear architecture and chromatin organization, potentially influencing gene cluster regulation implicated in pancreatitis pathogenesis. For instance, *Ehmt2* inactivation leads to derepressing the Beta Globin LCR, a regulatory region known for its involvement in internal chromosome looping. This potentially reflects reorganization of the 3D nucleus, suggesting a broader role for EHMT2 beyond direct transcriptional regulation and potentially influencing higher-order chromatin structures and nuclear organization. Emphasizing the interconnected nature of epigenetic regulation, transcriptional control, and nuclear organization in shaping inflammatory responses, future investigations into this phenomenon promise to offer further valuable insights into the molecular mechanisms underlying pancreatitis development.

In light of recent findings highlighting the enduring epigenetic memory of inflammatory injury in pancreatic acinar cells (Falvo et al. 2023), the current study’s investigations into *Ehmt2* knockout in this specific cell population during acute pancreatitis gains significance. At the core of our study lies a gene regulatory mechanism within pancreatic acinar cells, revealing how epigenetic dysregulation within this single cell type can instigate a cascade of events, ultimately leading to an amplified injury-inflammation-repair response across the entire organ. This discovery underscores the interconnectedness of cellular processes, where alterations in gene regulation within one cell type propagate across the organ’s cellular landscape, influencing its overall function and response to stimuli. Understanding these systemic effects will provide critical insights into organ homeostasis and disease progression. Moreover, shedding light on the interplay between EHMT2-mediated cell-specific epigenetic regulation and its broader effects on organ-level responses, our research provides a foundation for targeted interventions aimed at modulating EHMT2-related epigenetic memory and alleviating the long-term consequences of pancreatic injury.

Limited studies in acute pancreatitis have highlighted the involvement of epigenetic mechanisms, such as histone acetylation and methylation, as well as the therapeutic potential of targeting bromodomain and extra-terminal (BET) proteins and histone deacetylases (HDACs) in mitigating inflammation and reducing disease severity (Bombardo et al. 2017). Building upon this foundation, the insights gained from our study hold promise for the identification of EHMT2 as another potential therapeutic target, given its pivotal role in modulating inflammatory responses within the pancreas, thus adding to the growing list of epigenomic targets for intervention in pancreatic diseases. Strategies aimed at restoring EHMT2 function or inhibiting its downstream effectors could offer avenues for mitigating pancreatic inflammation and improving disease outcomes. Furthermore, the identification of EHMT2’s regulatory network provides a foundation for exploring combinatorial therapies that target multiple nodes within the inflammatory cascade. Such approaches may offer synergistic effects and enhance therapeutic efficacy. Overall, our study opens new avenues for therapeutic exploration in pancreatic diseases by elucidating the role of epigenetic regulators and their potential as targets for intervention.

In conclusion, our experiments in two complementary mouse models reveal the multifaceted role of EHMT2 in regulating gene expression and inflammation during acute pancreatitis, exacerbating pancreatic inflammation upon inactivation. This advances our understanding of EHMT2’s role in maintaining homeostasis and preventing severe inflammation. Importantly, these findings have implications beyond pancreatology, extending to inflammatory conditions in other organs, as recently evidenced by involvement of EHMT2 in autoimmune pathways, such as those observed in the colonic mucosa (Ramos et al. 2023). Additionally, the application of spatial transcriptomics further strengthens our results. Considering the ongoing evaluation and testing of EHMT2 inhibitors in preclinical studies in Sickle Cell Anemia, which also affect Beta Globin genes, and combination therapies for pancreatic cancer, our findings warrant careful consideration regarding their potential to attenuate pancreatic defenses against inflammatory stimuli (Krivega et al. 2015; Renneville et al. 2015; Katayama et al. 2020; Urrutia et al. 2020; Pan et al. 2016; Takase et al. 2023; Yuan et al. 2013). Overall, our study not only presents novel insights but also holds significant biomedical relevance in the context of autoimmune diseases, chronic inflammation, and emerging therapeutic strategies involving EHMT2.

## MATERIALS AND METHODS

### Mouse models and acute pancreatitis induction

Animal care and all experimental protocols were reviewed and approved by the Institutional Animal Care and Use Committees of Mayo Clinic Rochester (IACUC protocols A00002240-16 and A24815) and the Medical College of Wisconsin (AUA00005963). Mice were maintained in standard housing with controlled temperature, humidity, and light cycles and given standard rodent chow and water *ad libitum*. Tissues were collected and preserved in formaldehyde for a minimum of 24 hours prior to transfer to 70% (v/v) ethanol for histological processing and examination. *B6.FVB-Tg(Pdx1-Cre)6Tuv/J* (*Pdx1-Cre*, IMSR Cat# JAX:014647, RRID: IMSR_JAX:014647)(Hingorani et al. 2003) and *Ptf1a^TM^ ^1(cre)Hnak^/RschJ* (*P48^Cre/+^*, IMSR Cat# JAX:023329, RRID: IMSR_JAX:023329) (Nakhai et al. 2007) were originally purchased from Jackson Laboratories. *Ehmt2 flox/flox* (*Ehmt2^fl/fl^*) animals were generously provided by Dr. Oltz (Tachibana et al. 2007). Animals were maintained on a C57Bl/6 background, and genotyping procedures to confirm *Pdx1-Cre*;*Ehmt2^fl/fl^* and *P48^Cre/+^*;*Ehmt2^fl/fl^* crosses have been described previously (Urrutia et al. 2021). Both sexes were included in the experiments. Animals that were used in ontogeny studies with no additional treatments were sacrificed at 10 days (postnatal, PN) or 4 weeks (young adult, YA). For induction of acute pancreatitis, cohorts of 4-week-old mice were fasted for 12 hours before the first caerulein injection. Caerulein or saline control was administered via IP injection at a dose of 50 μg/kg once an hour for a total of 8 injections. Food was returned after the first dose. Animals were euthanized 18 hours after the initial dose. Tissue was taken (n=2-4) for RNA and histological analysis. Mice were euthanized using CO2 in accordance with institutional guidelines.

### Serum Analysis

Blood was collected from the animals by orbital puncture and serum was isolated for studying serum chemistry. Serum was mixed 1:1 with saline and their profile consisting of Glucose, Albumin, Globulin, Total Protein, Alanine Amino Transferase, Total bilirubin, Minerals (Sodium, calcium, Phosphorus) Blood Urea Nitrogen (BUN), and Alkaline phosphatase were measured using VetScan VS2 (Mathison et al. 2013).

### Histological analysis

Pancreatic tissues were paraffin-embedded for sectioning, and sections were stained with hematoxylin and eosin (H&E) for pathological evaluation. Pancreatitis severity was assessed by assessing pancreatic tissue edema, inflammatory cell infiltration, and necrosis. The scoring system was adapted from (Moreno et al. 2006). It included an assessment of pancreatic tissue edema (on a scale from a minimum of 0 for no edema to a maximum of 3 for separated and disrupted acini), inflammatory cell infiltrate (on a scale from a minimum of 0 for no infiltrate to a maximum of 3 with infiltrate in the parenchyma for >50% of the lobules), and necrosis (on a scale from a minimum of 0 for absence of necrosis to a maximum of 3 with diffuse parenchymal necrosis for >10% of the parenchyma). To quantify the severity of the parameters, four random fields (20× objective) per slide, each containing at least 1,000 cells per field, were imaged, and assessed by the three scales. Mean ± SD scores were calculated for each parameter.

### TUNEL assay

Formalin-fixed pancreatic tissues were paraffin-embedded and sectioned (5 µm). TUNEL analysis was carried out using the ApopTag Peroxidase in situ cell apoptosis detection kit (Millipore, S7100) according to the manufacturer’s directions. Slides were developed with Nova Red (Vector Laboratories) and counterstained with Mayer hematoxylin. Five random fields (20× objective) per section, containing at least 1,000 cells per field, were imaged and counted (Urrutia et al. 2020).

### RNA extraction, RNA-seq and Bioinformatics analysis

Preparation of RNA from tissue was performed as previously described (Urrutia et al. 2021). To reduce degradation during storage at −80°C, RNaseOUT (Invitrogen, Cat# 10777019) added to final elution. The RNA samples were quantified by Qubit (Invitrogen), and their quality was assessed using the Fragment Analyzer (Agilent). The samples with RINs > 6 and DV200 > 80% were selected for library preparation. The pancreas RNA was then sequenced using the Illumina TruSeq RNA v2 library preparation kit and the Illumina High Seq-2000. The sequencing reads were mapped to the mouse reference transcriptome Gencode vM23 (GRCm38.p6), and at least 24 million mapped read pairs were obtained per sample. The resulting reads were processed through the Mellowes Center workflow, which includes MapRseq3 (Kalari et al. 2014) and EdgeR (McCarthy, Chen, and Smyth 2012). Differential gene expression was based on a false discovery rate (FDR) < 0.1 and an absolute FC ≥ | 2.0|. Pathway analysis of DEGs was done using RITAN (Zimmermann et al. 2019) and the MSigDB hallmark gene set collection (Liberzon et al. 2015), while gene network and upstream regulatory analyses were performed using Ingenuity® Pathway Analysis (IPA®; Qiagen). To quantify immune-cell fractions in the young-adult/perinatal bulk RNA-seq comparisons, we performed digital cytometry analysis with the QuanTiSeq algorithms, which allowed intra-sample and inter-sample comparisons of only immune cell type fractions (Finotello et al. 2019). The quanTIseq method was applied on DEGs from through an R package called Immunedeconv allowing for identification of immune cell composition despite the low overall immune cell population (Sturm et al. 2019). Using the same R package for the bulk RNA-seq comparisons that involved caerulein injections, the MCP-Counter algorithm was used, which allowed for an expanded breakdown of the complete cell composition with robust quantification of the absolute abundance of eight immune and stromal cell populations (Becht et al. 2016).

### Spatial Transcriptomics

FFPE Pancreatic tissue sections (*Ehmt2^+/+^* and *Ehmt2^fl/fl^*) were deparaffinized followed by H&E staining and imaging using BZ-X800 (Keyence). Thereafter, tissue was destained, decrosslinked, and permeabilized for processing with the CytAssist and Visium Spatial Gene Expression Kits (10x Genomics; Pleasanton, CA, USA). Library quality metrics were confirmed by fragment analysis and Kapa qPCR before sequencing according to manufacturer recommendations on the NovaSeq 6000 (Illumina, San Diego, CA, USA). A sequencing depth of approximately 250–400 million read-pairs per sample was obtained. Sample processing, library preparation and sequencing for this project was completed by the Mellowes Center for Genomic Sciences and Precision Medicine Center at the Medical College of Wisconsin (RRID:SCR_022926). The original read quality was checked by FastQC and FastQ Screen. Alignment, tissue detection, fiducial detection, and barcode/UMI counting was performed by Spaceranger Count. For reads alignment, reference transcriptome is human GRCh38-2020-A with Visium Probe_Set_v2.0. Seurat was used for downstream spatial data analysis. First, spots with zero counts were filtered out, and normalization was performed using the SCTransform method. Subsequently, dimension reduction and clustering were performed with PCA and UMAP analysis. Spatially expressed genes for each cluster were found using the default Seurat method. Spatially variable genes were found using two Moran’s I methods, specifically the Seurat clustering and UMAP. Results were integrated to Loupe Browser for data visualization. Each cell type was assigned to its respective cluster through manual review of the expression of a comprehensive set of marker genes, as per outlined by (Cui Zhou et al. 2022), and the analysis of genes of interest involved utilizing the scale value, either in the form of Log normalized or Feature sum.

## ACKNOWLEDGEMENTS

This work was supported by NIH [grant numbers R01DK52913 (to R.U. and G.L.) and R01CA247898 (to G.L.)]; Advancing a Healthier Wisconsin Endowment (to G.L. and R.U.); the Linda T. and John A. Mellowes Endowed Innovation and Discovery Fund (to R.U.); The Joel and Arlene Lee Endowed Chair for Pancreatic Cancer Research (to G.L.); and Markus Family Funds for Discovery and Innovation Family Funds to Mellowes Center.

